# Expanded Analysis of the *Pantoea stewartii* subsp. *stewartii* DC283 Complete Genome Reveals Plasmid-borne Virulence Factors

**DOI:** 10.1101/407825

**Authors:** Duy An Duong, Ann M. Stevens, Roderick V. Jensen

## Abstract

*Pantoea stewartii* subsp. *stewartii*, a Gram-negative proteobacterium, causes Stewart’s wilt disease in corn. Bacterial transmission to plants occurs primarily via the corn flea beetle insect vector, which is native to North America. *P. stewartii* DC283 is the wild-type reference strain most used to study pathogenesis. Previously the complete genome of *P. stewartii* was released. Here, the method whereby the genome was assembled is described in greater detail. Data from a matepair library preparation with 3.5 kilobase insert size and high-throughput sequencing from the MiSeq Illumina platform, together with the available incomplete genome sequence of AHIE00000000.1 (containing 65 contigs) was used. This work resulted in the complete assembly of one circular chromosome, ten circular plasmids and one linear phage from *P. stewartii* DC283. A high number of sequences encoding repetitive transposases (> 400) were found in the complete genome. The separation of plasmids from genomic DNA revealed that two Type III secretion systems in *P. stewartii* DC283 are located on two separate mega-plasmids. Interestingly, the assembly identified a previously unknown 66-kb region in a location interior to a contig in the previous reference genome. Overall, a novel approach was successfully utilized to fully assemble a prokaryotic genome that contains large numbers of repetitive sequences and multiple plasmids, which resulted in some interesting biological findings.

## INTRODUCTION

*Pantoea stewartii* subsp. *stewartii* (referred to herein as *P. stewartii*), a Gram-negative gammaproteobacterium causes Stewart’s wilt disease in corn (1). *P. stewartii* belongs to the *Enterobacteriaceae* family containing plant-associate enterics (e.g. *Erwinia, Dickeya*, and *Pectobacterium* sp.) and human enteric and plant-associated pathogens (e.g. *Escherichia* and *Salmonella* sp.) (2). *P. stewartii* can colonize and grow to high cell density in the xylem of some varieties of corn, after being transmitted by the North American corn flea beetle, *Chaetocnema pulicaria* (3). The symptoms of Stewart’s wilt include water-soaked lesions when the infection occurs in the apoplast space at the early stages of the disease, wilting if the infection becomes systemic through the xylem vessels, and death if the plants were infected at their seedling phase (1, 4).

*P. stewartii* is a good model for studying bacterial quorum sensing cell-cell communication (5, 6), bacterial surface motility (7) and host-pathogen interactions (1, 7–14). *P. stewartii* interacts with both plant and insect hosts making its life-style interesting to investigate. There have been numerous studies focused on understanding the pathogenicity of this phytopathogen in order to counteract its effects on the plants and/or to break the infection cycle to protect the crop (1, 7–14). Two main virulence systems in *P. stewartii* have been characterized as the underlying mechanisms of wilt-disease symptoms. First, the Hrp-type III secretion system (T3SS) is linked to the water-soaked lesions that occur during the early stages of the disease (1, 12). Indeed, *P. stewartii* utilizes two T3SS to maintain its interaction in different hosts, one for corn and the other for the corn flea beetles (9, 15). Second, stewartan, a specific extracellular polysaccharide (EPS) produced by *P. stewartii* (16), is considered to be a main pathogenesis factor responsible for a number of disease symptoms such as vascular streaking, bacterial oozing and wilting (1, 17, 18). Stewartan production is controlled by quorum-sensing regulation at low cell density via the repression of RcsA, the transcription activator controlling EPS production (19). Quorum sensing is also known to regulate the surface motility of *P. stewartii,* which contributes to the virulence in the plants (7); however, the mechanism of this regulation remains unclear. Genes involved in pigment production (20), the regulation of oxidative stress (8), iron acquisition (14), the expression of an RTX-like cytolytic toxin (21), and the proteins Lon and OmpA (22) are additional *P. stewartii* virulence factors that have been discovered.

*P. stewartii* DC283, a nalidixic acid resistant mutant of the original *Zea mays* 1976 isolate SS104 (23, 24), is used as a wild-type reference strain to study pathogenesis of this phytopathogen. A draft genome assembly of *P. stewartii* DC283 was issued in 2012 with 65 contigs (NCBI GenBank: AHIE00000000.1). This enabled the initial genetic approaches to study this pathogen at larger scale, such as studying the whole transcriptome of the microorganism with RNA-Seq (25, 26). Since 2014, additional draft *Pantoea stewartii* genomes from different subspecies have been released but none were fully assembled (27). The difficulty of the whole genome assembly in *P. stewartii* DC283 could be explained in part by the constraints of the sequencing technologies available at the time, as well as the high number of plasmids naturally present in this bacterium (23) and the high amount of transposable elements in its genome (NCBI GenBank: AHIE00000000.1).

DNA sequencing has rapidly advanced with new technological developments over the last decade (28). Next-generation sequencing (NGS) now allows parallel sequencing of large numbers of relatively small fragments of DNA (reads) and assembly into large contiguous sequences (contigs) (29). Illumina is one of the four manufacturers providing currently available platforms to perform NGS (30). Illumina applies the optical detection of clonal amplification to determine the DNA sequence during the synthesis of the prepared templates (http://www.illumina.com). Mate-pair library preparation with large insertion size is a recent tool to improve the ability of the Illumina platform (31, 32) to complete the assembly of genomes, since it permits the extension and linkage of contigs terminated by long repetitive sequences (33). For this study 250 bp reads with a 3.5 kb insert were successfully used to generate linked contigs that were assembled into the complete genome of *P. stewartii* DC283. This promoted the analysis of essential genes *in planta* via Tn-Seq (22), facilitated a detailed analysis of the genome and revealed some distinctive features of interest in this genome.

## MATERIALS AND METHODS

### Library preparation and Illumina sequencing

*P. stewartii* DC283 was grown in 5 ml Luria Bertani (LB) medium (10 g/l tryptone, 5 g/l yeast extract, and 5 g/l NaCl) overnight and DNA was extracted using a QIAgen DNeasy Blood & Tissue Kit per the manufacturer’s recommendations for Gram-negative bacteria protocol. On-column RNase treatment was used prior to DNA elution of high-quality DNA for mate-pair library preparation and sequencing with paired-end sequencing technology (250 bp mate-pair reads with a 3,500 bp insert) on an Illumina MiSeq platform (Biocomplexity Institute, Virginia Tech, VA). The mate-pair library construction was performed using the Illumina Nextera Mate-Pair protocol, gel plus method. Genomic DNA was fragmented and selected for a desired size (~3.5 kb) on a Sage Science Pippin Prep using a 0.75% gel. These fragments were attached to labeled adapters at the two ends before circularization. Shearing of these circular constructs into smaller DNA pieces occurred on the Covaris m220 prior to an enrichment process for those fragments containing the two adapters with flanking DNA. Finally, the library was quantified using qPCR (Kapa Kit) and then pooled for sequencing on the MiSeq using a v2 500 cycle kit set to do 2X 250PE. The results of sequencing data were ~250 bp reads that can be traced back to their mates that were physically separated on the genomic DNA by ~3.5 kb.

### Bioinformatics analysis

Raw Illumina sequencing data was generated by the core sequencing facility in the form of fastq files. These fastq files were imported into Geneious V9.1.2 (Biomatters Ltd.) for further analysis. The mate-reads were first matched together using the ‘Set Paired reads’ function and then assembled using the ‘Map to Reference’ function to the 65 contigs from AHIE00000000.1 as the reference. Unmapped mate-pair reads were then *de novo* assembled to identify missing sequences and rearrangements of the reference contigs. The end sequences of the contigs were then extended ~3,500 bp by *de novo* assembling mate-pair reads where one mate maps to the last 3,500 bp of the contig. The Geneious *de novo* assembly was used on unmapped reads to identify missing sequences and link the gaps. Plasmids were identified and separated from the chromosomal DNA based on their average read coverage, as well as their ability to be circularized at two ends of their overlapping sequences.

### Assembly confirmation via PCR

Traditional PCR reactions were utilized to verify some of the connections between the contigs during the assembly process. In particular, PCR reactions were designed to verify the existence of the novel 66-kilobase (kb) sequence that resulted from the *de novo* assembly. Primers were designed to have a similar melting temperature (approximately 60°C) with a length between 20-30 bp in order to run them in parallel. Two pairs of forward and reverse primers were used to verify the connection between the 66-kb sequence with the interior of contig AHIE01000008 (PSG-1F/ PSG-1R & PSG-2F/ PSG-2R, Table 1). One*Taq*^®^ 2X Master Mix with standard buffer (New England Biolabs, USA) was used with a reaction volume of 15 μl and 667 nM of each primer. The thermocycler settings were 30 s at 94μC, 30 s at 55μC, and 1m 45s at 72μC for 30 cycles. Amplicons were visualized on a 1% agarose gel and then extracted prior to sequencing to confirm the generation of the proper products.

**Table 1.**
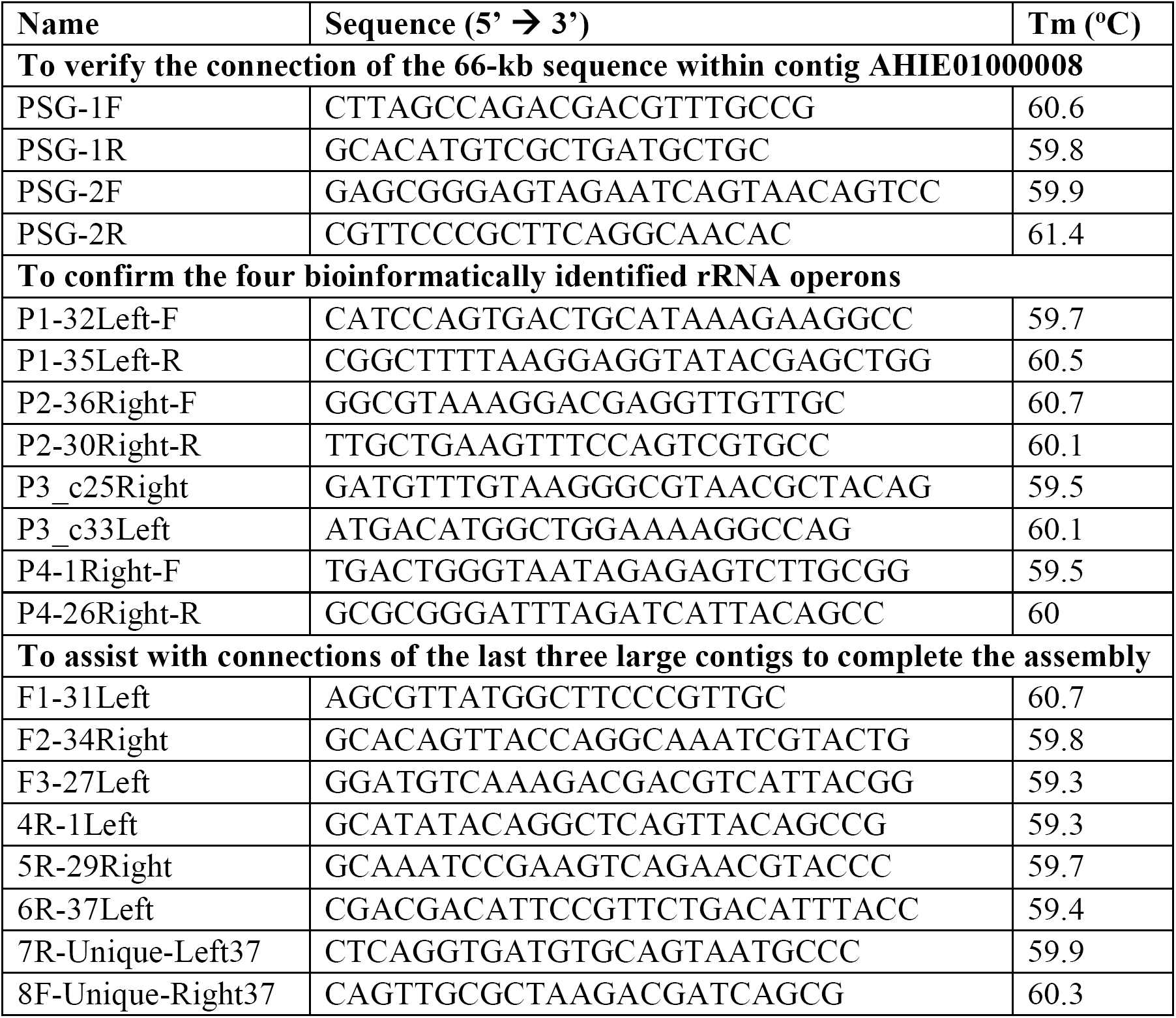
Primers used in this study

A QIAGEN^®^ LongRange PCR kit (Qiagen, USA) was used to amplify the four identified rRNA operons through bioinformatics analysis, which span approximately 6 kb, and to assist in the determination of the direction of connections of the last three large assembled chromosomal contigs whose ends contain multiple transposons, in addition one of these ends also had a rRNA operon. See Table 1 for the primer sequences, which were designed as described above. A reaction volume of 25 μl and 400 nM of each primer were used per the manufacturer’s instruction. The thermocycler settings were 15 s at 93μC, 30 s at 55μC, and 7 m at 68oC for 35 cycles. Amplicons were visualized on a 1% agarose gel and extracted prior to sequencing.

### Nucleotide sequence accession number

As previously published in *Genome Announcements* (34), the complete genome was annotated using the NCBI prokaryotic annotation pipeline and the complete genome assembly of *P. stewartii* DC283 is available in GenBank under accession no. CP017581-CP017592.

### Development of PCR method to screen for the presence of the 66-kb region

A multiplex PCR reaction with three primers for rapid detection of the missing 66-kb region was developed for the analysis of new *P. stewartii* strain constructs. This reaction contained three primers PSG-1F, PSG-2F and PSG-2R at 667 nM concentration each in a 15 μl final volume using One*Taq*^®^ 2X Master Mix with standard buffer (NEB). The thermocycler settings were 30 s at 94μC, 30 s at 55μC, and 1 m 45 s at 72μC for 30 cycles. The amplicon was visualized on a 1% agarose gel to determine its size. This corresponds to the presence (1736 bp product of PSG-2F and PSG-2R) or absence (1324 bp product of PSG-1F and PSG-2R) of the 66-kb region. The reaction was then repeated with PSG-1F and PSG-1R to confirm the presence or absence of the 66-kb sequence.

## RESULTS

### Comparison of the previous incomplete and new complete genome sequences of *P. Stewartii* DC283

The complete genome of *P. stewartii* DC283 consists of 5,314,092 bp (53.8% G+C content) including 5,625 coding sequences, 73 tRNAs and 21 rRNAs (39). This genome includes one circular chromosome, ten circular plasmids and one linear phage (Table 2). The previous genome assembly (AHIE00000000.1) consisting of 65 contigs was complicated by the large number of repetitive transposon sequences that prematurely terminated contigs and introduced ambiguities in the assembly. Our complete assembly identified 444 repetitive transposon sequences often spanning ~1,500 bp that could nevertheless be bridged using the mate-pair reads with 3,500 bp inserts. The mate-pair reads linked the ends of the previous 65 contigs and led to rearrangements of the sequences in some of them. Table 3 summarizes some major features of the sequencing results for the complete *P. stewartii* DC283 genome generated in this study in comparison with the initial incomplete reference sequence (AHIE00000000.1).

**Table 2.** DNA components in the complete assembly of the *P. stewartii* DC283 genome.

| **Name** | **Length (bp)** | **Topology** | **Mean Coverage** | **Copy Number** |
| --- | --- | --- | --- | --- |
| Chromosome | 4,528,215 | circular | 181.28 | 1 |
| pDSJ01 | 4,277 | circular | 3355.36 | 21 |
| pDSJ02 | 4,368 | circular | 2214.90 | 14 |
| pDSJ03 | 13,414 | circular | 1414.80 | 9 |
| pDSJ04 | 25,681 | circular | 952.56 | 6 |
| pDSJ05 | 34,447 | circular | 593.62 | 4 |
| pDSJ06 | 47,481 | circular | 414.55 | 3 |
| pDSJ07 | 65,483 | circular | 408.61 | 2 |
| pDSJ08 | 105,961 | circular | 440.82 | 3 |
| pDSJ09 | 132,938 | circular | 277.89 | 2 |
| pDSJ010 | 304,641 | circular | 163.45 | 1 |
| ppDSJ01 | 47,186 | linear | 963.65 | 6 |

**Table 3.** Comparison between the incomplete (AHIE00000000.1) and the complete (CP017581-CP017592) assembly of *P. stewartii* DC283 genome

| Feature | AHIE000000000.1 | CP017581-CP017592 |
| --- | --- | --- |
| Total length (bp) | 5,233,214 | 5,314,092 |
| G+C % | 53.8 | 53.8 |
| Total DNA content | 65 contigs | 1 chromosome, 10 plasmids,<br>1 linear phage plasmid |
| # Proteins | 4,903 | 4,942 |
| # Pseudogenes | 164 | 411 |
| # tRNA genes | 70 | 73 |
| # rRNA genes | 20 | 21 |
| # genes annotated as encoding a transposase | 361 | 444 |

### Two type III secretion systems in *P. stewartii* are located on two separate mega-plasmids

The main circular chromosome was assembled using multiple rounds of mate-pair recognition and mapping, together with long-range PCR, to rearrange and connect most of the contigs from the incomplete reference sequence (AHIE00000000.1). The same approach was used to connect and circularize the remaining contigs of the incomplete genome into the 10 circular plasmid sequences. Several of these sequences were highly similar to known *P. stewartii* plasmids sequences, e.g. the closure of AHIE01000052 was found to be similar to pSW100 (35) and is now known as pDSJ01 (4,277 bp), the closure of AHIE01000062 was found to be similar to pSW200 (36) and is now known as pDSJ02 (4,368 bp), and the closure of AHIE01000065 was found to be similar to pSW800 (37) and is now known as pDSJ05 (34,447 bp). Preliminary annotation was performed by rapid annotation using subsystem technology (RAST, http://rast.nmpdr.org/). This approach was used to identify genes involving in the replication of plasmids (eg. *repA*) in the circular sequences prior to submission to NCBI for final annotation. In addition, the amount of read coverage determined during assembly was used to calculate the average copy number of each plasmid in *P. stewartii* DC283. Ten separate plasmids with their copy numbers were identified and renamed in order from smallest to largest according to their molecular mass (Table 2). The small plasmids pDSJ01 and pDSJ02 exist as intermediate-level copy numbers in *P. stewartii* while the other plasmids and the linear phage-plasmid were present at lower copies per cell. The three megaplasmids (size above 100 kb) have the lowest copy number, from 1-3 copies per cell.

The separation of the plasmids from the genomic DNA and their annotation revealed that the two T3SS in *P. stewartii* DC283 are located on two separate mega-plasmids. Specifically, genes related to the T3SS needed for the invasion of the insect host and colonization of the plant host (9) are located in plasmid pDSJ08 and plasmid pDSJ10, respectively, in the complete genome.

### Identification of a phage N15-like linear phage plasmid of *P. stewartii*

There are a high number of sequence reads mapped to contig AHIE01000047 (46,532 bp) in the draft genome having a coverage of ~1000X, which may correspond to approximately six copies of this sequence in the *P. stewartii* DC283 genome. This contig also contains genes encoding a partitioning system usually found in plasmid sequences. However, the contig end extension using mate-pair reads did not result in a connection with any other contigs or self-circularization. This suggests that the sequence is a linear extrachromosomal DNA element present in multiple copies. The annotation of this contig also showed a high number of phage-related coding sequences. In addition, data from the blastx function on the NCBI website against nonredundant protein sequences (nr) database for this sequence showed some level of similarity to coding sequences in a N15 prophage in *Escherichia coli* (38). Therefore, this contig-end fixed sequence has been renamed as ppDSJ01 (47,186 bp) to represent a linear phage plasmid of *P. stewartii* DC283 (Table 2).

### Multiple chromosomal prophage sequences found in *P. stewartii* genome

When the sequenced reads were mapped to the draft contig AHIE01000008, an interesting mapping coverage was observed (Figure 1) from position ~140,000 to ~200,000. The average coverage of the entire contig is ~320X with the majority of the sequence having coverage at ~200X which is the baseline of coverage of the chromosomal DNA as a single copy in a cell. However, the specific region of DNA shown in Figure 1 has three regions of elevated coverage. Two of these regions have around two times the coverage of the chromosomal DNA and a region of up to ~16 times the chromosome coverage in the middle region spanning between positions of ~165,000 and ~180,000 of contig AHIE01000008. The entire elevated region spans from position ~2,032,000 to ~2,089,000 of the complete *P. stewartii* DC283 chromosome (CP017581) with the middle region spanning between positions of ~2,053,000 and ~2,066,000. The annotation of these elevated coverage regions showed that they contain genes related to bacteriophage sequences. It was not possible to distinguish whether these sequences belong to tandem sequence duplications in the chromosome in this region or whether some of them are also present in the cell as extrachromosomal linear phage DNAs.

**Figure 1.**
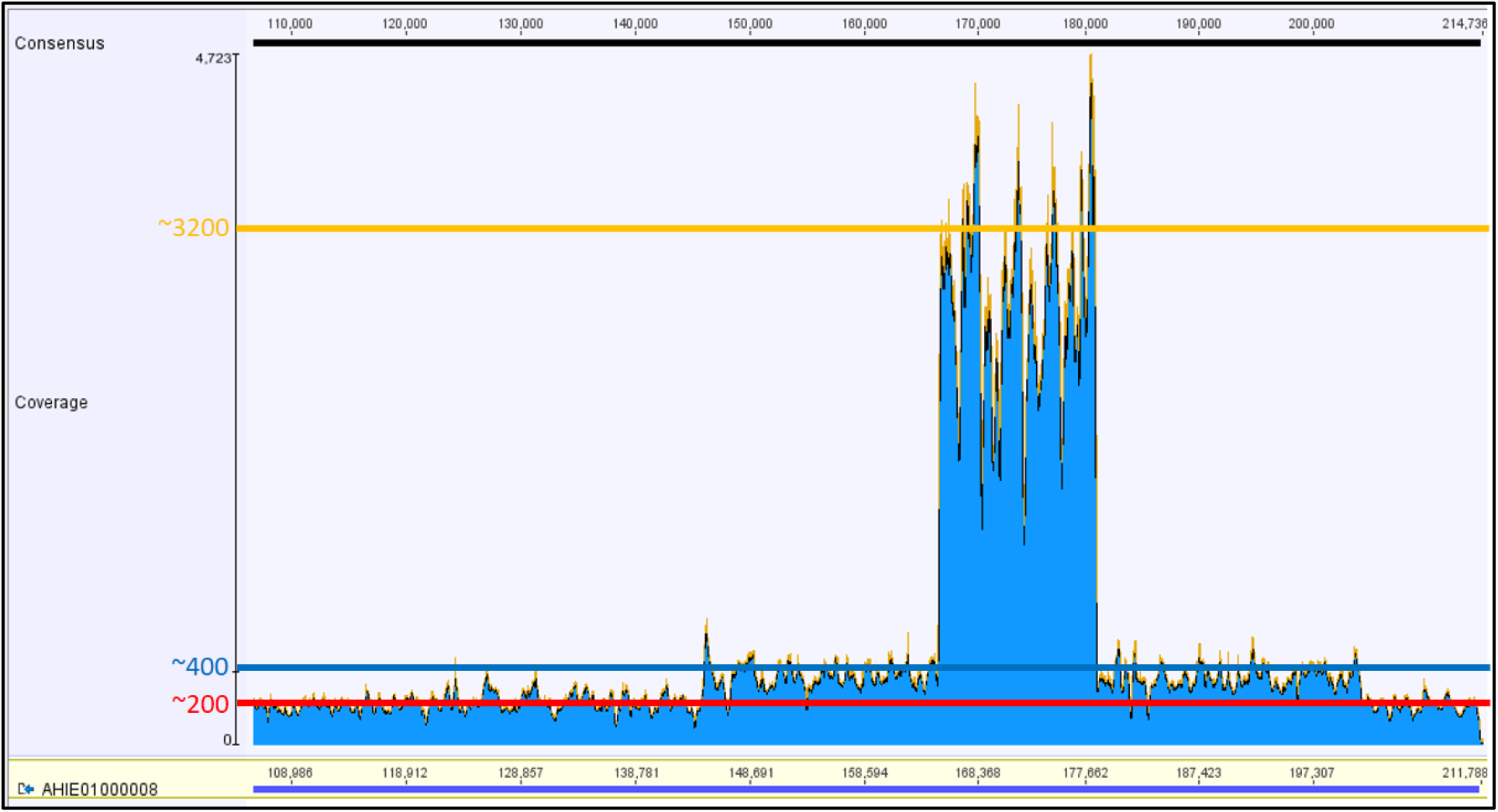
Coverage of mapped reads to a region on contig AHIE01000008 of the incomplete genome. The average coverage of chromosomal DNA is ~200X (the red line) while two regions, spanning ~140,000 -~165,000 and ~180,000 -~200,000 of contig AHIE01000008, show an average of ~400X (the blue line). The yellow line indicates another region, spanning ~165,000 -~180,000 of contig AHIE01000008, with a coverage at ~3200X. The elevation in coverage suggests either tandem duplication of the sequence or the presence of the same regions extrachromosomally in one or multiple copies.

Similarly, another region spanning from ~2,411,000 to ~2,463,000 of the complete *P. stewartii* chromosome DNA molecule was recognized as duplication in the sequence coverage suggesting either tandem repeat of the ~51,000 bp region or the extrachromosomal existence of a linear DNA molecule with the same sequence chromosomally integrated. The annotation of genes included inside this region showed a large amount of hypothetical proteins and phage-related proteins.

### Seven contigs in the reference genome are absent in the complete *P. stewartii* genome

Surprisingly, there are seven small contigs in the incomplete genome with no sequence reads mapped to their sequences on the new complete genome. The total length of these contigs is 23,196 bp with 17 genes (Table 4). This content of DNA makes up only 0.4 % of the total DNA length of the new assembly of the *P. stewartii* DC283 genome. It could not be determined whether these contigs are real and reflect a discrepancy between the two DNA genome sequences of *P. stewartii* DC283, or they are results of sequencing and assembly errors. However, the annotation of the genes located on these contigs suggested that they may be part of extrachromosomal DNA such as plasmids or mobile elements like transposons (Table 4) in the previous draft genome, producing genetic variation between strains.

**Table 4.**
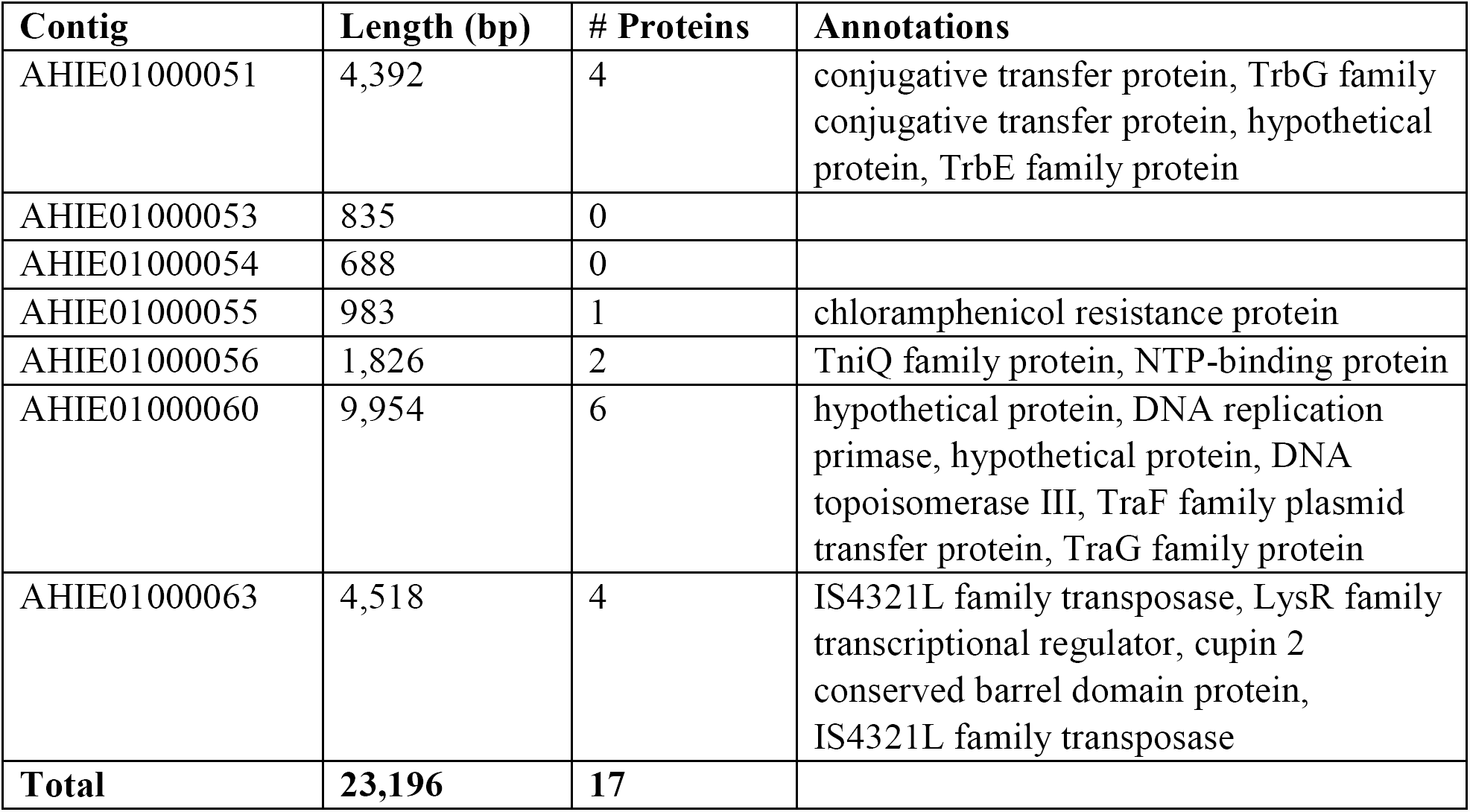
Seven contigs present in the incomplete genome but absent in the complete genome of *P. stewartii* DC283

| Contig | Length (bp) | # Proteins | Annotations |
| --- | --- | --- | --- |
| AHIE01000051 | 4,392 | 4 | conjugative transfer protein, TrbG family<br>conjugative transfer protein, hypothetical<br>protein, TrbE family protein |
| AHIE01000053 | 835 | 0 |  |
| AHIE01000054 | 688 | 0 |  |
| AHIE01000055 | 983 | 1 | chloramphenicol resistance protein |
| AHIE01000056 | 1,826 | 2 | TniQ family protein, NTP-binding protein |
| AHIE01000060 | 9,954 | 6 | hypothetical protein, DNA replication<br>primase, hypothetical protein, DNA<br>topoisomerase III, TraF family plasmid<br>transfer protein, TraG family protein |
| AHIE01000063 | 4,518 | 4 | IS4321L family transposase, LysR family<br>transcriptional regulator, cupin 2<br>conserved barrel domain protein,<br>IS4321L family transposase |
| **Total** | **23,196** | **17** |  |

### Identification of a novel region of *P. stewartii* genome

Interestingly, the assembly of the new sequencing data resulted in the identification of a novel region, 66 kb in length containing 68 genes, in the interior of contig AHIE01000008 of the previous reference genome. PCR reactions were used to confirm the existence of this region inside this contig (Figure 2). These PCR results followed by Sanger sequencing of the PCR products have confirmed the integrity of the bioinformatics analysis using *de novo* assembly of the unmapped reads and the contig end extension methods. This region is now located from 1,862,325 to 1,928,315 of the *P. stewartii* main chromosome (CP017581). A multiplex PCR reaction with primers designed to confirm the connection of this region to the interior of contig AHIE01000008 was successfully used to screen for the presence or absence of this 66-kb region in *P. stewartii* strains. Only one previously published deletion strain and its complement (25) did not have the newly identified 66-kb region whereas this region is present in the rest of the tested strains.

**Figure 2.**
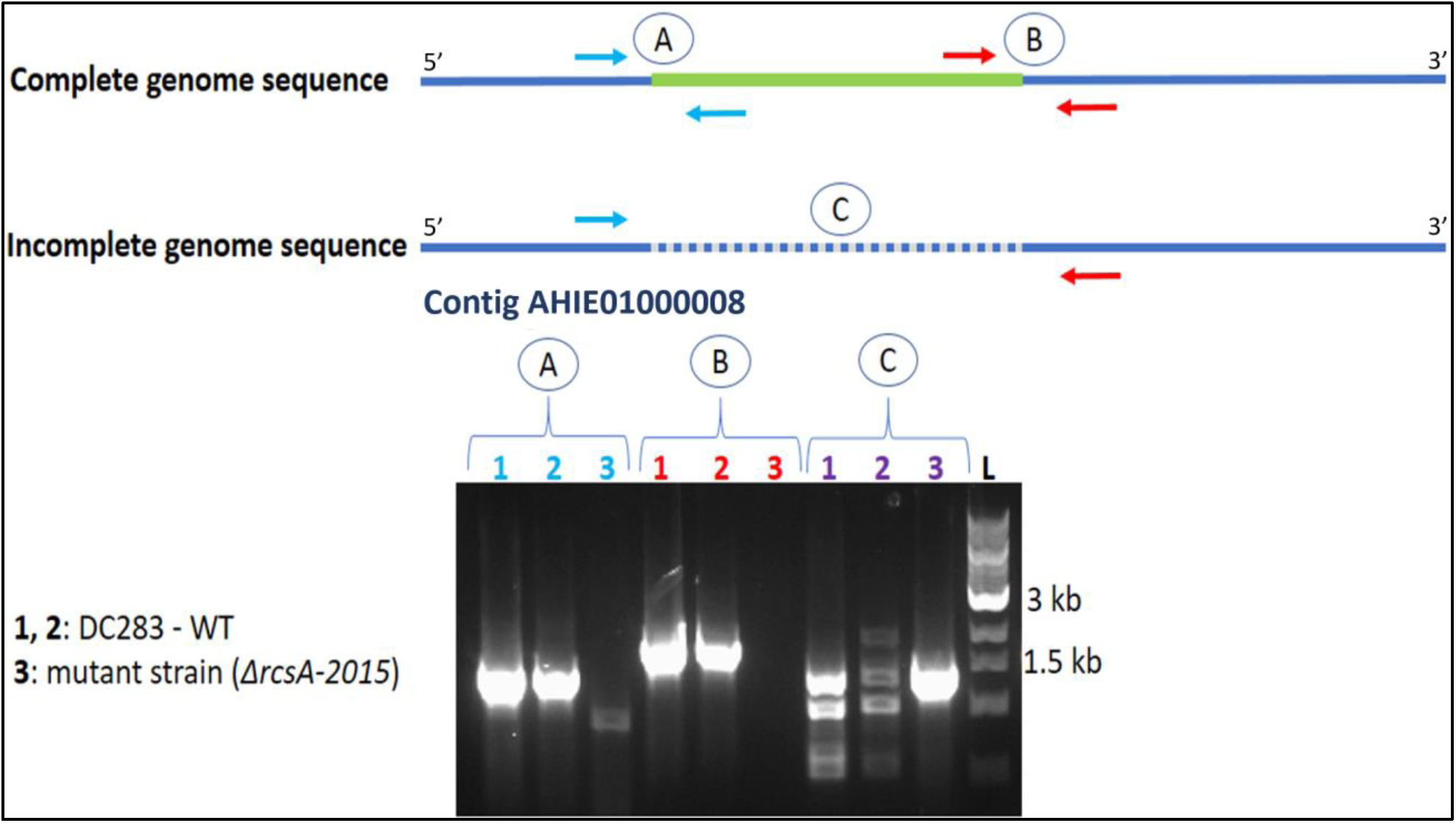
Confirmation of the identification of the *de novo* assembled 66-kb region using traditional PCR reactions. Each PCR reaction contains two primers to detect the presence of the physical connection in the tested strains. Two wild-type (WT) isolates and one mutant (*ΔrcsA2015*) strain, were used to demonstrate the presence/absence of the appropriate specific PCR product. A: connection between the 5’ end of the 66-kb region (green line) to the 3’ end of the interior of contig AHIE01000008; B: connection between the 3’ end of the 66-kb region (green line) to the 5’ end of the interior of contig AHIE01000008; C: a region interior of contig AHIE01000008 in the incomplete genome sequence (AHIE00000000.1). In A and B, only wildtype strains show the specific products indicating the presence of this 66-kb region. In C, the outer primers demonstrate the continuity of this sequence in the mutant strain with a specific PCR product in comparison with multiple unspecific products present in the wild-type samples.

## DISCUSSION

A novel approach involving mate-pair library preparation with paired-end Illumina sequencing technology was successfully utilized to fully assemble the *P. stewartii* DC283 bacterial genome that contains large numbers of repetitive sequences and multiple plasmids. This whole assembly of the *P. stewartii* genome has revealed a rich content of extrachromosomal DNA. Two small plasmids, each ~4.5 kb, were present at a medium copy number whereas the other plasmids are maintained in less than 10 copies per cell. The larger the size of a plasmid, the less abundant it seems to be in *P. stewartii* cells. This negative correlation between plasmid size and their copy number in a bacterial cell was previously experimentally determined in *Bacillus thuringiensis* YBT-1520 (39). The genome of *Bacillus thuringiensis* YBT-1520 contains 11 plasmids (between 2 kb and 416 kb in size) with their copy numbers negatively correlated to their molecular mass. This general trend may correspond to the energetic burden of carrying a plasmid and maintaining its copy number inside the cells (40, 41).

The two T3SS used by *P. stewartii* to independently colonize two different hosts and maintain its life cycle between the plants and the insect vectors (9) are now known to be maintained on two mega-plasmids (pDSJ010 and pDSJ008), respectively. The largest *P. stewartii* plasmid, pDSJ010, contains the plant colonizing T3SS and it is the universal plasmid LPP-1 in the genus *Pantoea* (42). LPP-1 plasmids, derived from an ancestral plasmid, contain a large repertoire of proteins that contribute to the adaptation of species of the genus to their various niches and their specialization as beneficial biocontrol agents, harmless saprophytes, or pathogens (42). Other plasmids carried by *P. stewartii* may also play important roles in many cellular biophysical processes as well as colonization and pathogenesis within multiple hosts. The complete sequence for these plasmids will facilitate the utilization of bioinformatic approaches to understand their roles in relation to the host bacterium.

This project also resulted in the identification of the presence of an extrachromosomal linear plasmid phage in multiple copies which is similar to the lambdoid phage N15 of *Escherichia coli* (38). Additional phage elements found in the *P. stewartii* genome at positions ~2,032,000 ~2,089,000 and ~2,411,000 ~2,463,000 could be intrachromosomally incorporated in tandem and/or present as linear plasmid phages with several copies per cell. Further investigation is needed to clarify this issue in the assembly as well as to understand the role of phage elements inside this bacterial host.

Interestingly, a small amount of DNA sequence (~23 kb) from the incomplete genome was not present in the newly assembled genome of *P. stewartii* DC283. This might have arisen through technical discrepancies between the methods used to generate the two genome sequences. Alternatively, these missing sequences may belong to unstable DNA molecules that led to genetic content differences among the reference strains of *P. stewartii* DC283 after they were domesticated in multiple laboratories for decades. The second reason seems to be probable for at least some of these sequences when the annotation is taken into account. The majority of these genes code for conjugative transfer components of plasmids, phage-related proteins, and transposon elements (Table 4).

The recognition of the 66-kb region missing in the reference genome and a mutation strain (*rscA* deletion strain (25)) generated from our laboratory led to two findings. The first one is the existence of this region in the genome which may be involved in several different metabolic pathways since it contains 68 coding sequences. This also raised a concern about the stability of the genetic material within this species. A rapid PCR detection method was developed in this study to screen for the presence of this region among genetically modified strains (43). This preliminary survey resulted in the confirmation of the presence of this region in all test strains with the exception of the *rcsA* deletion strain. This provides confidence in the overall stability of this region among *P. stewartii* laboratory strains. However, the frequency of elimination of this region, what triggers it and the consequential biological effects of this region on the bacteria during infection remain unknown.

In conclusion, this project has provided a detailed analysis of the complete genome of *P. stewartii*. This phytopathogen has a complex genome containing multiple plasmids, phages and other mobile elements such as transposons. The complete genome has facilitated further understanding of this pathogen, as well as the *Pantoea* genus, using bioinformatic analysis and high-throughput sequencing technology.

## ACKNOWLEDGEMENTS

We thank Susanne von Bodman and David Mackey for *P. stewartii* strains. This work was funded by the Fralin Life Science Institute (RJV) and Biological Sciences Department (AMS), Virginia Tech. Mate-pair library construction and Illumina sequencing were conducted by Genomics Research Laboratory at the Biocomplexity Institute at Virginia Tech.

